# Tissue-specific vulnerability to apoptosis in Machado-Joseph disease

**DOI:** 10.1101/2023.04.09.536069

**Authors:** Ana F. Ferreira, Mafalda Raposo, Emily D. Shaw, Naila S. Ashraf, Filipa Medeiros, Maria de Fátima Brilhante, Matthew Perkins, João Vasconcelos, Teresa Kay, Maria do Carmo Costa, Manuela Lima

## Abstract

Machado-Joseph disease (MJD) is a dominant neurodegenerative disease caused by an expanded CAG repeat in the *ATXN3* gene encoding the ataxin-3 protein. Several cellular processes, including transcription and apoptosis, are disrupted in MJD. To gain further insights into the extent of dysregulation of mitochondrial apoptosis in MJD, and to evaluate if expression alterations of specific apoptosis genes/proteins could be used as transcriptional biomarkers of disease, the levels of *BCL2, BAX* and *TP53* and the *BCL2*/*BAX* ratio, an indicator of susceptibility to apoptosis, were assessed in blood and *post-mortem* brain samples from MJD subjects and MJD transgenic mice and controls. While patients show reduced levels of blood *BCL2* transcripts, this measurement displays low accuracy to discriminate patients from matched controls. However, increased levels of blood *BAX* transcripts and decreased *BCL2*/*BAX* ratio are associated with earlier onset, indicating a possible association with MJD pathogenesis. *Post-mortem* MJD brains show increased *BCL2*/*BAX* transcript ratio in the dentate cerebellar nucleus (DCN) and increased BCL2/BAX insoluble protein ratio in the DCN and pons, suggesting that in these regions, severely affected by degeneration in MJD, cells show signs of apoptosis resistance. Interestingly, a follow-up study of 18 patients further shows that blood *BCL2* and *TP53* transcript levels increase over time in MJD patients. Furthermore, while the similar levels of blood *BCL2, BAX*, and *TP53* transcripts observed in preclinical subjects and controls is mimicked by pre-symptomatic MJD mice, the expression profile of these genes in patient brains is partially replicated by symptomatic MJD mice. Globally, our findings indicate that there is tissue-specific vulnerability to apoptosis in MJD subjects and that this tissue dependent behavior is partially replicated in a MJD mouse model.

## 1. Introduction

Machado-Joseph disease/spinocerebellar ataxia type 3 (MJD/SCA3) (MIM#109150) is the most frequent autosomal dominant inherited ataxia worldwide [1]. MJD is caused by an expansion of a polyglutamine (polyQ)-encoding CAG repeat in the *ATXN3* gene [2]. Ataxin-3 (ATXN3), the protein encoded by *ATXN3*, is ubiquitously expressed in different cell types of peripheral and brain tissues [3]. Neuropathological complexity is a characteristic of this disease, which affects especially the cerebellum, brainstem, basal ganglia, some cranial nerves and spinal cord [4]. In average the age at onset of MJD has been reported around the 40 years-old, and the mean survival time is estimated to be of 21 years [5]. In MJD, overt disease is preceded by a preclinical phase in which neuropathological and molecular alterations are already present [6–9].

Several studies in cellular and animal models of MJD, including the YACMJDQ84.2 (Q84) transgenic mice that is frequently used in pre-clinical studies of MJD [10–14], have been revealing that mutant *ATXN3* harboring an expanded polyQ tract is involved in the disruption of transcription, impairment of mitochondrial function, alteration of mechanisms regulating oxidative stress, as well as apoptosis [3,15,16]. Several of these changes, such as transcriptional dysregulation, have also been observed in blood samples from MJD carriers [17,18].

Mitochondrial apoptosis is a complex mechanism regulated by several proteins including those belonging to the BCL2 family, such as the anti-apoptotic BCL2 (B-cell lymphoma-2) and the pro-apoptotic BAX (BCL2 Associated X) [19]. Another crucial player in apoptosis is the transcription factor P53 (Tumor protein p53) which, among other roles, regulates the levels and the activity of BCL2 family proteins [20]. In fact, evidence of a direct link between mutant ATXN3 and apoptosis has been provided by Liu and colleagues [21] who reported that p53 is a substrate of ATXN3 and that mutant ATXN3, exhibiting a stronger deubiquitinase activity than native ATXN3, increases p53 stability and its consequent accumulation in MJD cellular models, leading to increased p53-dependent apoptosis in a MJD zebrafish model and in HCT116 cells. Moreover, our group has formerly shown that peripheral blood cells of MJD carriers display decreased *BCL2*/*BAX* ratio [8] resulting from a reduction of blood *BCL2* transcript levels [18], indicating a higher vulnerability to an apoptotic stimulus in MJD subjects compared with controls. Similarly, at the protein level, Bcl2 levels and Bcl2/Bax ratio have been found to be decreased in human neuronal SK-N-SH cells expressing expanded-poly-Q ATXN3 compared with the parental cells [22]. Although some studies using animal and cellular models have shown the involvement of neuronal apoptosis in MJD [22,23], little is known about the expression patterns of apoptotic-related genes in the brain and other tissues of individuals with MJD [24].

Hence, the levels of the apoptosis-related *BCL2, BAX* and *TP53* transcripts in cross-sectional and longitudinal samples of peripheral blood from MJD carriers (patients and pre-clinical subjects) were assessed to gather further insights into the dysregulation of mitochondrial apoptosis in MJD and to evaluate their capacity of being used as transcriptional biomarkers of MJD. Additionally, the expression patterns of *BCL2, BAX* and *TP53* genes were investigated in samples from *post-mortem* brains of MJD patients to better elucidate the involvement of the mitochondrial apoptosis pathway in MJD pathogenesis and to evaluate possible conserved changes of apoptosis markers in the brain and peripheral blood of MJD patients. Finally, the transcript and protein levels of mouse *Bcl2, Bax* and *Trp53* genes were assessed in samples from the peripheral blood and the brain of Q84 transgenic mice, expressing the full-length human *ATXN3* gene harboring an expanded CAG repeat, to determine whether this widely used MJD mouse model replicates the findings observed in MJD subjects.

## 2. Materials and Methods

### 2.1. Human samples and clinical data

A total of 19 preclinical (no ataxia or diplopia) subjects, 42 MJD patients and 63 healthy controls were included in this study. A summarized clinical, genetic, and demographic characterization of the individuals included in this study is provided in Supplementary Table S1.

Blood samples: baseline samples from 19 preclinical subjects, 37 patients and 54 age (±3 years) and sex-matched paired controls were used in a cross-sectional study to evaluate the expression levels of *BCL2, BAX* and *TP53* transcripts (Figure 1, A1; Supplementary Table S1). A subset of 13 patients with five or less years of disease duration was used to represent the early stage of the disease (early patients) (Figure 1, A1). The cut-off used to define the early patients was based on the natural history data for MJD that shows that the mean score of disease progression (defined by gait impairment) evolves very slowly during the first five years [25]. Additionally, for a subset of 18 MJD patients, blood samples collected at one or two additional observational moments were used in a follow-up study (Figure 1, A1; Supplementary Table S2). All MJD subjects underwent a standard neurological evaluation by the same single neurologist. Age at onset was defined as the age of appearance of the first symptoms (gait ataxia and/or diplopia) reported by the patient or a close relative. Disease duration was defined as the time elapsed between age at onset and age at neurological evaluation/blood collection. Preclinical subjects were enrolled in the study after being confirmed as carriers of the *ATXN3* mutation, in the context of the Azorean Genetic Counseling and Predictive Test Program offered by the regional health system to patients and families. For preclinical subjects the number of years between the age at neurological evaluation/blood collection and the predicted age at disease onset (years to onset, time) was calculated as previously described by Raposo and colleagues [26]. The determination of the number of CAG repeats at the *ATXN3* locus was performed for all MJD subjects as described by Bettencourt and colleagues [27]. In addition to the lack of family history of MJD, the majority of the controls were molecularly excluded for the *ATXN3* mutation using the above-mentioned protocol [27]. The use of blood samples and clinical data of MJD carriers for the purposes of this study was approved by the Ethics Committee of the University of the Azores (Parecer 5/2017 and Parecer 26/2018). All subjects provided written informed consent for research.

**Figure 1.**
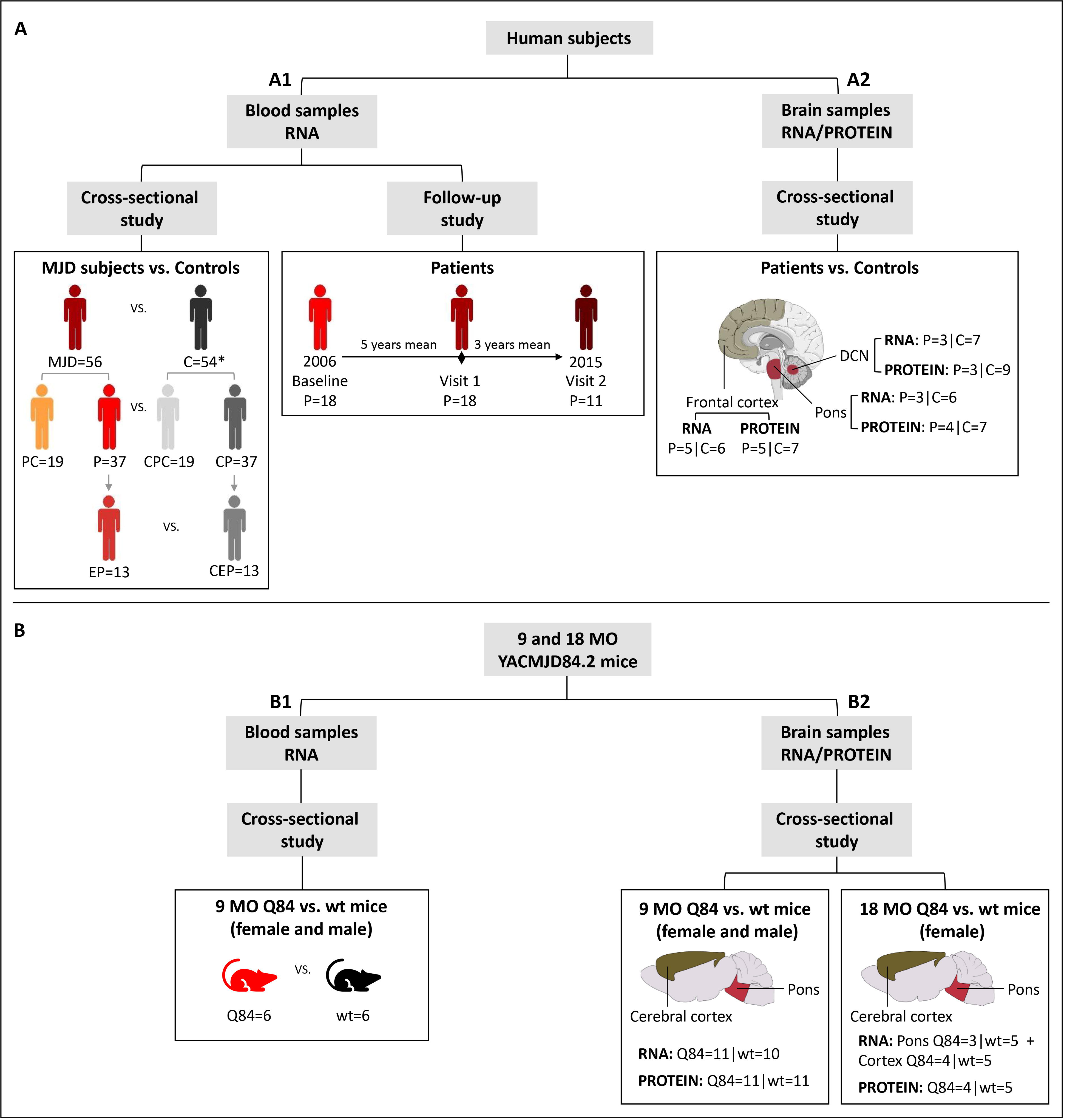
Design of the study: BCL2, BAX and TP53 transcript and protein levels were assessed in (A) human subjects and in the (B) YACMJD84.2 (Q84) mouse model. In human subjects, the expression levels were analyzed in (A1) blood samples from MJD carriers (preclinical subjects and patients) and controls in a cross-sectional and from patients in a follow-up study, and in (A2) brain samples from MJD patients and controls in a cross-sectional study (brain figure was partly generated using Servier Medical Art, provided by Servier, licensed under a Creative Commons Attribution 3.0 unported license). Expression levels of the mouse homologue Bcl2, Bax, and Trp53 genes were analyzed in (B1) blood samples from 9-month-old (MO) Q84 transgenic and wild-type (wt) littermate mice and in (B2) brain samples from 9 and 18 MO Q84 transgenic and wt mice. MJD, MJD subjects; PC, preclinical subjects; P, patients; EP, early patients; C, controls; CPC, controls of PC subjects; CP, controls of patients; CEP, controls of early patients. *two control individuals were included in both CPC and CP groups.

Brain samples: frozen samples from *post-mortem* brain regions of deidentified and molecularly confirmed MJD patients (n=5) and molecularly excluded control subjects (n=9) were obtained from the University of Michigan Brain Bank (Table 1; Supplementary Table S3) and were used in a cross-sectional study (Figure 1, A2). Samples from two brain regions severely affected in MJD (dentate cerebellar nucleus (DCN) and pons) [4] and from a region with lower degree of disease affectation (frontal cortex) [28,29], were used in this study. The *ATXN3* CAG repeat number of all *post-mortem* brain samples was determined at Laragen Inc. (Culver City, CA) by DNA fragment analysis (Supplementary Table S3). The use of *post-mortem* brain samples for the purposes of this study was approved by the University of Michigan Biosafety Committee (IBCA00000265) and by the Ethics Committees of the University of the Azores (Parecer 26/2018).

### 2.2. Mouse samples

Hemizygous YACMJD84.2 (Q84) and wild-type (wt) littermate mice [30] were generated and maintained at the University of Michigan in a C57BL/6J background strain [10,14]. Mice were housed in cages with a maximum of five animals (range 3-5) and maintained in a standard 12h light/dark cycle with food and water ad libitum. Hemizygous Q84 mice show a reduced locomotor and exploratory activity on an open-field test at least at 75 weeks of age (≈17 months-old-MO) [10]. At least at eight weeks of age, hemizygous Q84 transgenic mice show some neuropathological features of MJD compared to non-transgenic littermates, such as progressive intranuclear accumulation of ATXN3 in neurons of several brain regions that are known to be also affected in MJD patients [10]. Pre-symptomatic 9 months-old (MO) Q84 (n=11) and wt mice (n=11), and symptomatic 18 MO Q84 (n=4) and wt mice (n=5) were used in the present study (Figure 1, B; Supplementary Table S4). Mouse genotyping for the presence of the human disease *ATXN3* transgene was performed using DNA isolated from tail biopsy at the time of weaning, as previously described by Costa and colleagues [10], and confirmed using DNA extracted from the tail collected *post-mortem*. The mouse *ATXN3* CAG repeat size was determined as above mentioned for human brain samples. Animal procedures were approved by the University of Michigan Committee on Use and Care of Animals (Protocol PRO00006371). The use of animal samples for the purposes of this study was approved by the Ethics Committees of the University of the Azores (Parecer 26/2018).

Blood samples: mice were anesthetized with ketamine/xylazine, blood was collected from a subset of the 9 MO mice (Q84=6 and wt=6) (Figure 1, B1) with a sterile syringe by cardiac puncture, and RNAlater Stabilization Solution (Thermo Fisher Scientific) was added to the blood and stored at −20°C. Mice were subsequently perfused transcardially with phosphate-buffered saline.

Brain samples: 9 MO (Q84=11 and wt=11) and 18 MO (Q84=4 and wt=5) mouse brains (Figure 1, B2) were collected after saline perfusion, dissected in the two hemispheres, and pons and cerebral cortex (isocortex) regions were macro dissected and stored at −80°C. Brain tissues from the right hemisphere were used for RNA studies and tissues from the left hemisphere were used for protein studies.

### 2.3. RNA extraction and cDNA synthesis

Human samples: total RNA was extracted from blood samples according to the procedures described in Raposo and colleagues [17]. cDNA was generated from 0.5µg of total RNA using the High-Capacity cDNA Reverse Transcription Kit (Applied Biosystems). For *post-mortem* brain samples, total RNA was extracted using Trizol (Invitrogen), and purified using the RNeasy mini kit (Qiagen) as previously described [10]. RNA concentration and integrity were assessed using the Agilent 2100 Bioanalyzer (Supplementary Table S3). cDNA was generated from 1.25µg of total RNA using the iScript cDNA synthesis kit (BIORAD).

Mouse samples: total RNA from mouse blood samples was extracted using the RiboPure RNA Purification Kit (Invitrogen). Total RNA from brain regions was extracted, quantified, and evaluated for integrity as described above for *post-mortem* human brain samples.

### 2.4. Quantitative real-time PCR

Human samples: qPCR experiments were conducted using the SensiFAST Probe Hi-ROX Master Mix (Bioline) and the TaqMan Gene Expression Assays (Applied Biosystems) for *BCL2* (Hs00608023_m1), *BAX* (Hs00180269_m1) and *TP53* (Hs01034249_m1) using an ABI StepOnePlus Real-Time PCR System (Applied Biosystems). Each sample was run in triplicate (blood) or quadruplicate (brain) alongside with the respective reference gene in the same plate. Experiments including the subset of patients used in the follow-up study were performed by including the samples from a given subject corresponding to all available collection moments in the same plate. In qPCR experiments using RNA from blood samples and *post-mortem* brain samples, *TRAP1* (TNF receptor associated protein 1; Hs00972326_m1) [31] and *GAPDH* (glyceraldehyde 3-phosphate dehydrogenase; Hs02758991_g1) were respectively used as reference genes. The relative expression values were calculated using the 2 method [32] in DataAssist v3.0 software (Applied Biosystems).

Mouse samples: the qPCR experiments using mouse blood and brain samples were conducted as above described for human blood samples, using the following TaqMan Gene Expression Assays (Applied Biosystems) for *Bcl2* (B cell leukemia/lymphoma 2; Mm00477631_m1), *Bax* (BCL2-associated X protein; Mm00432051_m1) and *Trp53* (transformation related protein 53; Mm01731287_m1). For both blood and brain samples, *Hmbs* (hydroxymethylbilane synthase; Mm01143545_m1) was used as the reference gene.

### 2.5. Western Blotting

Human samples: protein extracts from DCN, pons and frontal cortex tissues were obtained by homogenization in cold phosphate-buffered saline (PBS) containing protease inhibitor cocktail (cOmplete Protease Inhibitor Cocktail, Roche) and phosphatase inhibitors (PhosSTOP, Roche), followed by sonication and centrifugation. The supernatant was collected (PBS-soluble fraction) and stored at −80°C. The pellet was resuspended in 1% sodium lauroyl sarcosinate (sarkosyl, Sark)/PBS, sonicated, centrifuged and the supernatant (sarkosyl-soluble fraction) was stored at −80°C. Total protein concentrations from PBS-soluble (soluble) and sarkosyl-soluble (insoluble) fractions were assessed using the BCA method (Pierce BCA Protein Assay Kit; Thermo Fisher Scientific). Total protein lysates from PBS-soluble fractions and sarkosyl-soluble fractions (50µg) were resolved on 12% SDS-PAGE gels, and corresponding polyvinylidene difluoride (PVDF) membranes were incubated overnight at 4°C with various antibodies: rabbit anti-Bcl-2 (1:500; #3498, Cell Signaling), rabbit anti-Bax (1:500; #2772, Cell Signaling,) mouse anti-p53 (1:1000; #2524, Cell Signaling), mouse anti-GAPDH (1:10000; MAB374, Millipore), and mouse anti-ATXN3 (1H9) (1:2000; MAB5360, Millipore). Bound primary antibodies were visualized by incubation with peroxidase affiniPure goat anti-rabbit or anti-mouse secondary antibody (1:10000; Jackson Immuno Research Laboratories), followed by reaction with ECL-plus reagent (Western Lighting, PerkinElmer) and subsequent exposure to autoradiography films. Film band intensity was quantified by densitometry on ImageJ.

Mouse samples: protein extraction and quantification from pons and cerebral cortex from 9 and 18 MO Q84 and wt mice was conducted as previously described for *post-mortem* human brain samples. Total protein lysates from PBS-soluble and sarkosyl-soluble fractions (40µg 75µg) of each brain region were resolved on 12% SDS-PAGE gels.

### 2.6. Statistical analysis

#### 2.6.1. Cross-sectional studies

Human samples: blood transcript levels of MJD subjects were compared to those of controls using a T-test. When considering the groups of preclinical subjects and patients individually, blood transcript levels were compared to age- and sex-matched paired controls using a Wilcoxon Signed-rank test (two-tailed). Sex of preclinical subjects and patients was compared using a Chi-square test, and age at blood collection and the expanded CAG repeat size were compared using a Mann-Whitney U test (two-tailed). The ability for blood transcript levels to discriminate MJD subjects from healthy controls was assessed using the receiver operating characteristics (ROC) curve analysis; ROC curve accuracy was measured by the area under de curve (AUC) (95% confidence interval (CI), p-value). Correlations between blood transcript levels and clinical features of MJD carriers were conducted using the Spearmańs rank correlation (two-tailed); a partial correlation (two-tailed) was used to adjust for covariates, whenever necessary. *Post-mortem* brain transcripts and protein levels were compared between biological groups (patients and controls) using a generalized linear model adjusting to the age at death.

Mouse samples: blood and brain transcript and protein levels of Q84 and wt mice (9MO males and females combined) were compared using a Mann-Whitney U test (two-tailed).

GraphPad Prism version 8.0.1 for Windows (GraphPad Software, San Diego California USA) was used to identify extreme outliers (ROUT method Q=1%), which were excluded from data analysis, and to generate the data cross-sectional graphs. Statistical analysis was performed using IBM SPSS Statistics for Windows, version 25 (IBM Corp., Armonk, N.Y., USA). Statistical significance was set at p<0.05.

#### 2.6.2. Follow-up study of human blood samples

A total of 18 MJD patients were included in the follow-up study. Since repeated observations were performed on the same subjects over a period of 9 years (2006 to 2015), a linear mixed model was considered. The time metric used in the level 1 model was the baseline year 2006. The information about age at blood collection, sex, expanded CAG repeat length, age at onset and disease duration were used in the level 2 model, to determine the relationship between blood transcript levels of *BCL2, BAX* and *TP53* as well as *BCL2*/*BAX* ratio and the above-mentioned variables. The statistical analysis was performed using R software building on the longitudinal data analysis described by Garcia and colleagues [33]. Statistical significance was set at p<0.05.

## 3. Results

### 3.1. Human subjects

#### 3.1.1. Reduced transcript levels of the anti-apoptotic *BCL2* gene in blood of MJD patients

A cross-sectional study was conducted to evaluate if the transcript levels of *BCL2, BAX* and *TP53* are dysregulated in peripheral blood samples from total MJD subjects (Figure 2A) or in samples from subgroups of preclinical subjects, patients, and early patients, comparing with age- and sex-matched paired controls (Figure 2B-C). While comparison of patients with matched controls suggests reduced *BCL2* levels in MJD patients (p=0.041) (Figure 2B), *BCL2* transcript abundance displays low accuracy to discriminate patients from matched controls (AUC=0.64 (0.51-0.77), p=0.030).

**Figure 2.**
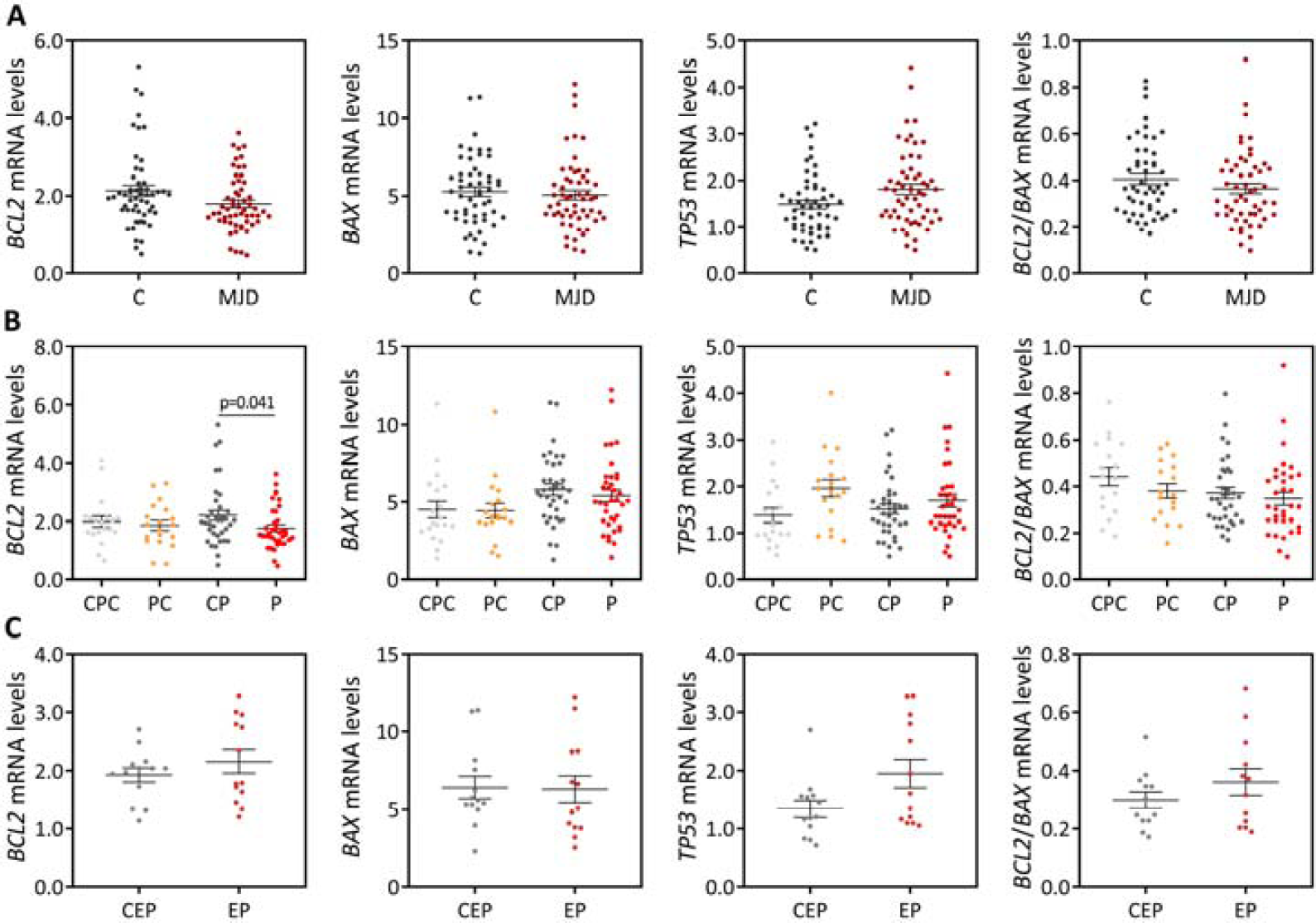
BCL2, BAX and TP53 transcript levels and BCL2/BAX ratio in peripheral blood samples from MJD carriers and controls. (A) Overall MJD subjects were compared with matched controls. (B) Preclinical subjects (PC) and patients (P) were compared to age- and sex-matched paired controls (CPC and CP, respectively). (C) Early patients (EP) were compared with age- and sex-matched paired controls (CEP). Graphs show the mean and standard error mean of the 2 values.

#### 3.1.2. Blood samples of MJD patients with earlier onset display higher levels of the pro-apoptotic *BAX* and lower *BCL2*/*BAX* ratio

Blood transcript levels of *BCL2, BAX* and *TP53* as well as the *BCL2*/*BAX* ratio were correlated with MJD subjects’ demographic, genetic and clinical features (Supplementary Table S5). After adjusting for the age at blood collection and the number of CAG repeats in the expanded *ATXN3* allele, higher transcript levels of *BAX* and lower *BCL2*/*BAX* ratio were observed in MJD patients with earlier onset (rho=-0.482, p=0.003 and rho=0.393, p=0.022, respectively) (Supplementary Table S5).

#### 3.1.3. Blood anti-apoptotic *BCL2* and pro-apoptotic *TP53* transcript levels increase over time in MJD patients

*BCL2, BAX* and *TP53* blood transcript levels and *BCL2*/*BAX* ratio alterations during disease progression were evaluated in a subset of 18 MJD patients for whom at least two observation points were available (Supplementary Figure S1). When adjusting a linear mixed model to the expression data of each of the three genes and to the *BCL2*/*BAX* ratio values, *TP53* levels increased on average 0.117 per year of disease (95% CI=0.063-0.172, p=0.001). Although in a lower extent, *BCL2* levels also increased on average 0.049 per year of disease (95% CI=0.001-0.098, p=0.046). Noteworthy, none of the possible confounding variables (age at blood collection, sex, expanded CAG repeat, age at disease onset and disease duration) influenced the observed increase of *TP53* and *BCL2* transcript abundance over time.

#### 3.1.4. *BCL2*/*BAX* ratio is increased in the degenerative DCN and pons of MJD patients

The expression behavior of *BCL2, BAX, TP53* was next assessed at the transcript and protein levels in a small set of *post-mortem* brain samples from MJD patients and control individuals. Three brain regions of MJD patients were evaluated in this study: DCN and pons which are greatly affected by degeneration in MJD patients [4], and the less affected frontal cortex [28,29]. Comparisons of transcript levels between MJD patients and controls for each brain region solely showed an increase of *BCL2*/*BAX* ratio in the DCN of patients (p=0.005) (Figure 3).

**Figure 3.**
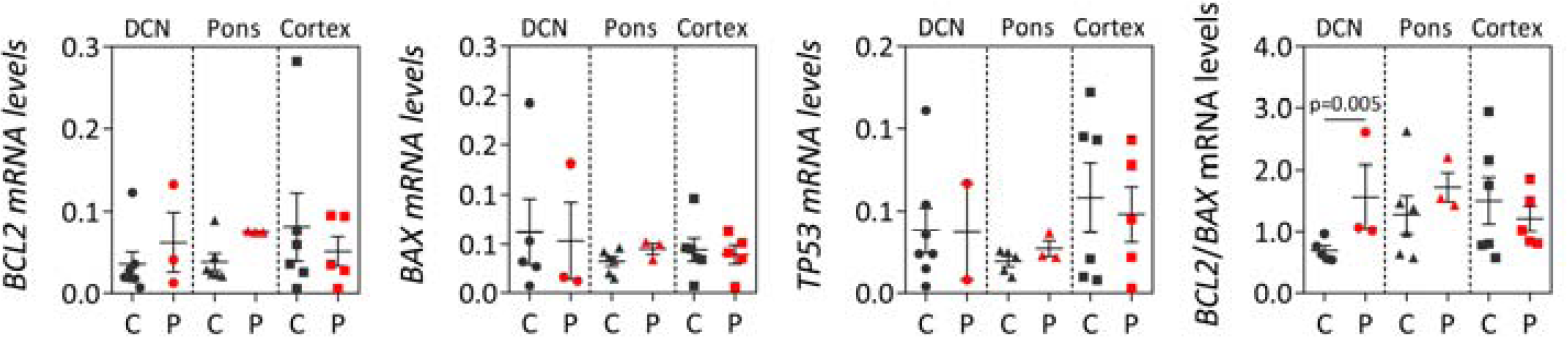
BCL2, BAX and TP53 transcript levels and BCL2/BAX ratio in post-mortem brain regions (dentate cerebellar nucleus (DCN), pons and frontal cortex) from MJD patients (P) and controls (C). Graphs show the mean and standard error of the mean of 2 values.

Next, protein levels of BCL2, BAX and P53 were analyzed both in the soluble fraction (enriched in cytoplasmic proteins) and in the insoluble fraction (enriched in nuclear and insoluble proteins, including membrane proteins) of brain lysates (Figure 4; Supplementary Figure S2). In DCN, compared with controls, patients showed increased BCL2/BAX ratio in the insoluble fraction (p=0.003) (Figure 4A). In pons, comparison between patients and controls revealed that patients show reduced soluble BCL2 levels (p=0.004) and subsequently decreased soluble BCL2/BAX ratio (p=0.028), and higher insoluble BCL2 levels (p=0.005) and consequently increased of insoluble BCL2/BAX ratio (p<0.001) (Figure 4B). In the frontal cortex, compared to controls, patients only showed increased soluble BAX levels (p=0.039) (Figure 4C). Importantly, BCL2/BAX ratio was increased in patients DCN both at the transcript and insoluble protein levels compared with healthy controls.

**Figure 4.**
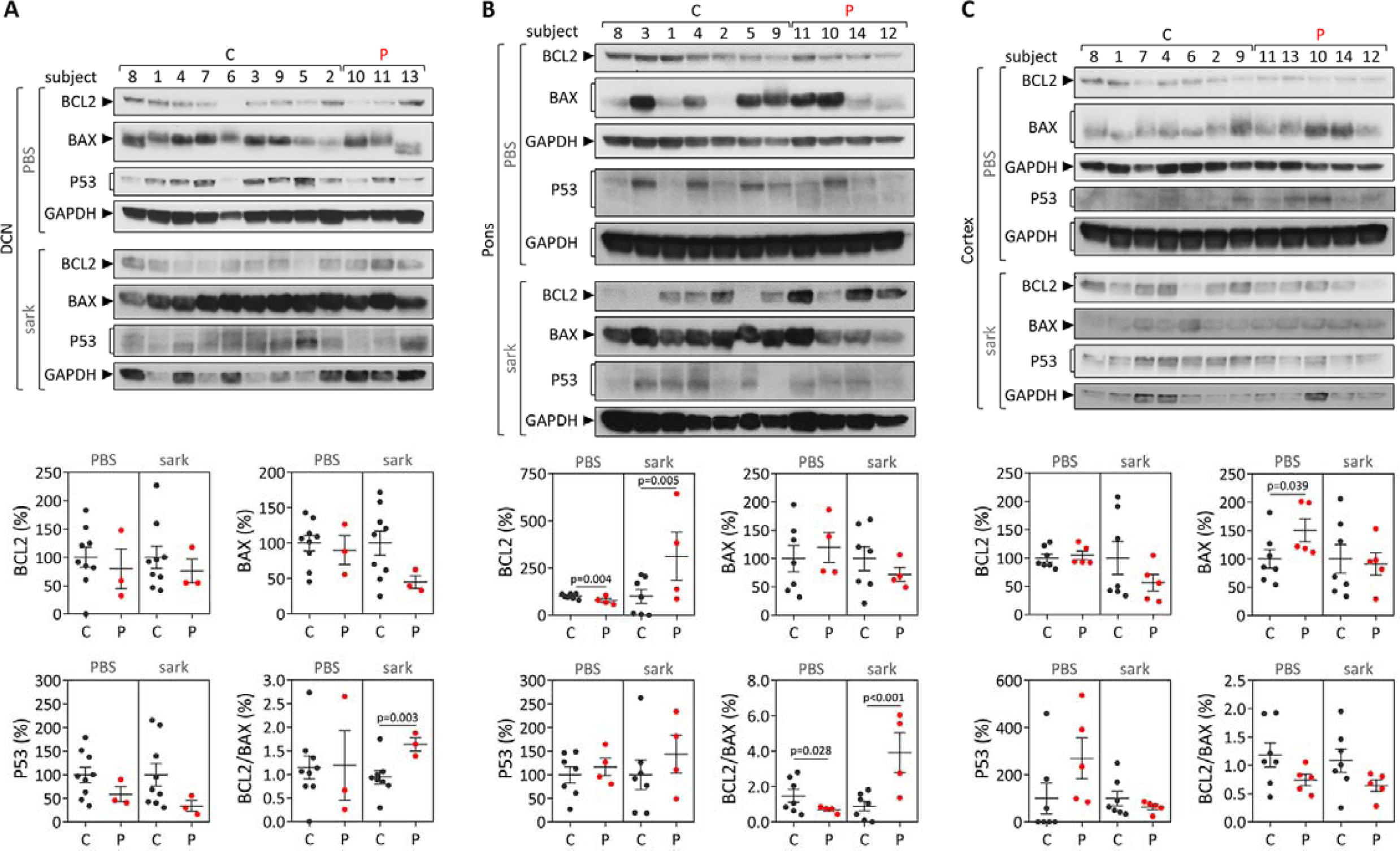
BCL2, BAX and P53 protein levels and BCL2/BAX ratio in post-mortem brains from MJD patients (P) and controls (C). Protein levels in PBS-soluble (soluble) and sarkosyl-soluble (insoluble) fractions were assessed in (A) dentate cerebellar nucleus (DCN), (B) pons and (C) frontal cortex. Upper panels show immunoblots of indicated proteins in both soluble (PBS) and insoluble (sark) fractions. Lower panels display quantification of band intensity, with values normalized to GAPDH. Bars in the graphs represent the average percentage of protein relative to controls (± SEM.

### 3.2. MJD mouse model

#### 3.2.1. Abundance of *Bcl2, Bax* and *Trp53* transcripts is similar in blood and brain samples of Q84 and wt mice

To determine whether Q84 mice replicate the blood transcriptional behavior of *BCL2, BAX* and *TP53* of preclinical subjects, the levels of their homologue genes were analyzed in blood samples of pre-symptomatic 9MO Q84 mice (without motor dysfunction [10]) and wt littermates. Similar to the observed when comparing preclinical MJD subjects with matched controls, no significant differences of *Bcl2, Bax* and *Trp53* blood transcript levels or Bcl2/Bax ratio were found in pre-symptomatic 9 MO Q84 mice compared to wt mice (Figure 5A). Next, the levels of the three genes were evaluated in the pons and cerebral cortex, respectively, affected and mildly/non-affected regions by degeneration [30], of pre-symptomatic 9 MO and symptomatic 18 MO Q84 mice [10] and wt littermate controls. As previously found in MJD patients’ pons and frontal cortex samples, no differences of transcriptional levels were observed for the three evaluated genes in the two corresponding brain regions of Q84 mice (Figure 5B-C).

**Figure 5.**
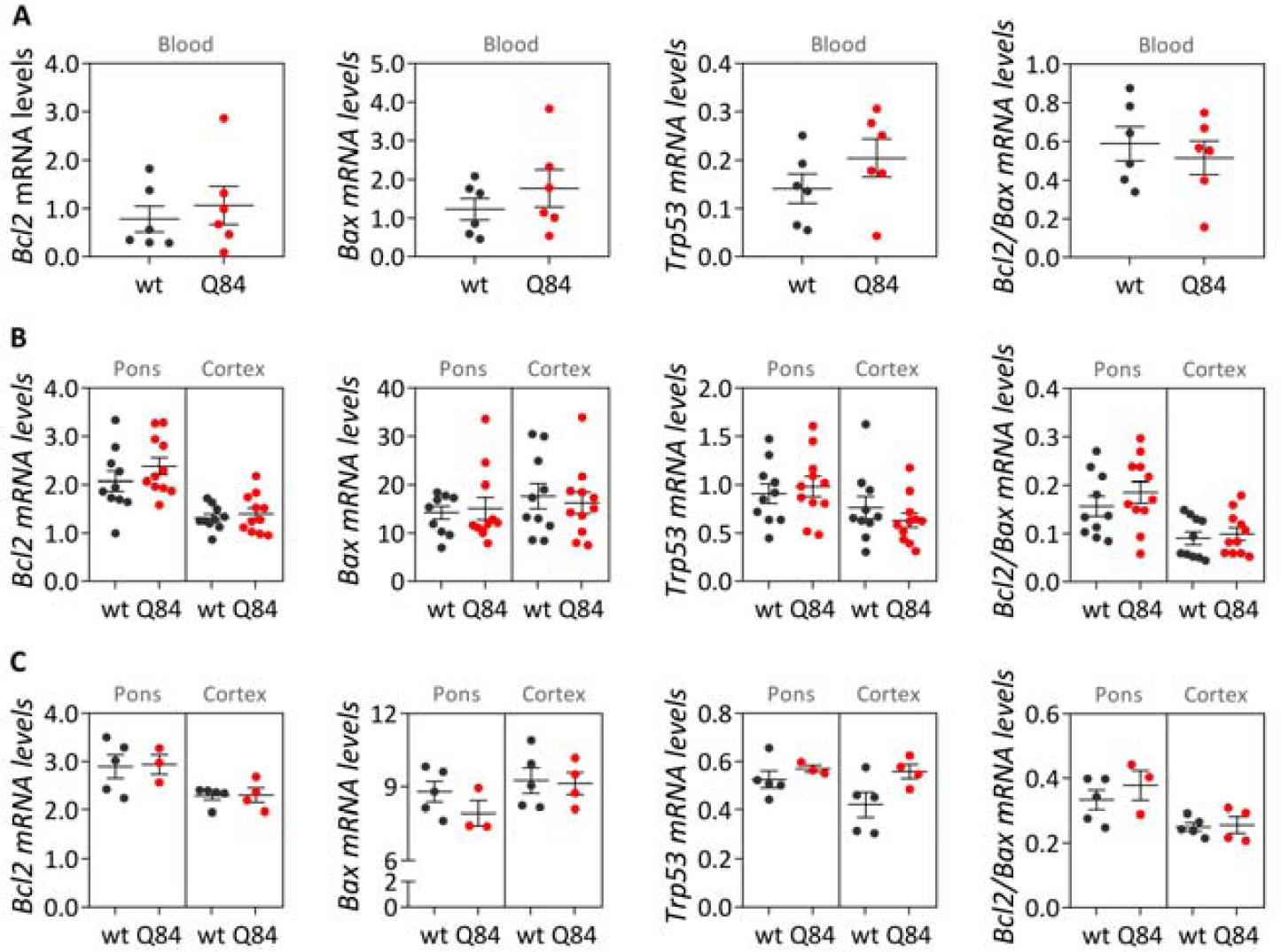
Transcript levels of Bcl2, Bax and Trp53 and Bcl2/Bax ratio in pre-symptomatic 9 months-old (MO) (A-B) and symptomatic 18 MO (C) Q84 transgenic and wt mice. Transcript levels were analyzed in (A) blood samples and in (B) brain samples from pons and cerebral cortex of the same mice. Graphs show the mean and standard error of the mean of 2 values.

#### 3.2.2. The pons and the cerebral cortex of pre-symptomatic Q84 mice show evidence of increased abundance of soluble BAX protein but similar Bcl2/Bax ratios comparing with wt mice

Protein levels of Bcl2, Bax and p53 were analyzed in the soluble and insoluble fractions of brain lysates from pons and cerebral cortex of pre-symptomatic 9 MO Q84 transgenic [10] and wt mice (Figure 6; Supplementary Figure S3). In pons, comparing with wt mice, Q84 mice suggests increased soluble Bax levels (p=0.001) and increased insoluble Bcl2 levels (p=0.003) (Figure 6A). In cerebral cortex, Q84 mice suggests increased soluble Bax levels (p=0.033) (Figure 6B).

**Figure 6.**
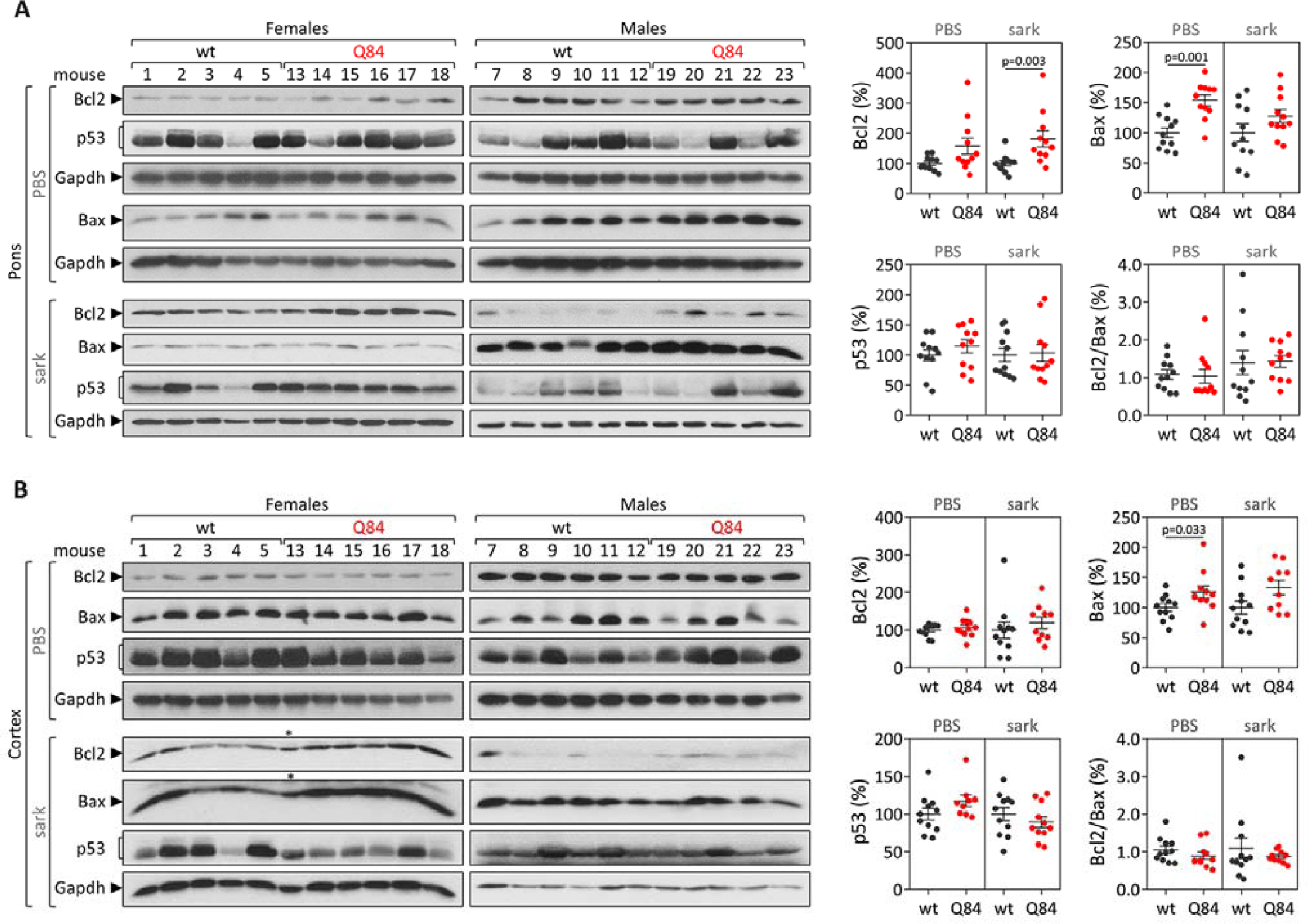
Protein levels of Bcl2, Bax and p53 as well as Bcl2/Bax ratio in PBS-soluble (soluble) and sarkosyl-soluble (insoluble) fractions were assessed in (A) pons and (B) cerebral cortex of pre-symptomatic 9 MO Q84 transgenic and wt mice. Left panels show immunoblots of indicated proteins in both soluble (PBS) and insoluble (sark) fractions. Right panels display the quantification of band intensity of all samples, with values normalized to GAPDH. Bars in the graphs represent the average percentage of protein relative to controls (± SEM).

#### 3.2.3. Cerebral cortex from symptomatic Q84 transgenic mice shows evidence of higher insoluble Bcl2/Bax ratio

Protein levels of Bcl2, Bax and p53 were further analyzed in the soluble and the insoluble fractions of pons and cerebral cortex lysates from symptomatic 18 MO Q84 transgenic (showing motor impairment [10] and wt mice. In pons, comparison between symptomatic Q84 and wt mice suggests reduced soluble Bax levels (p=0.014) and a subsequent increase of soluble Bcl2/Bax ratio in Q84 transgenic mice (p=0.027) (Figure 7A). In cerebral cortex, symptomatic Q84 mice solely suggests increased insoluble Bcl2/Bax ratio compared with wt mice (p=0.027) (Figure 7B). These results contrast with the findings in the corresponding brain regions from the MJD patients (decreased of soluble Bcl2/Bax ratio in the pons and increased insoluble Bcl2/Bax ratio in the frontal cortex).

**Figure 7.**
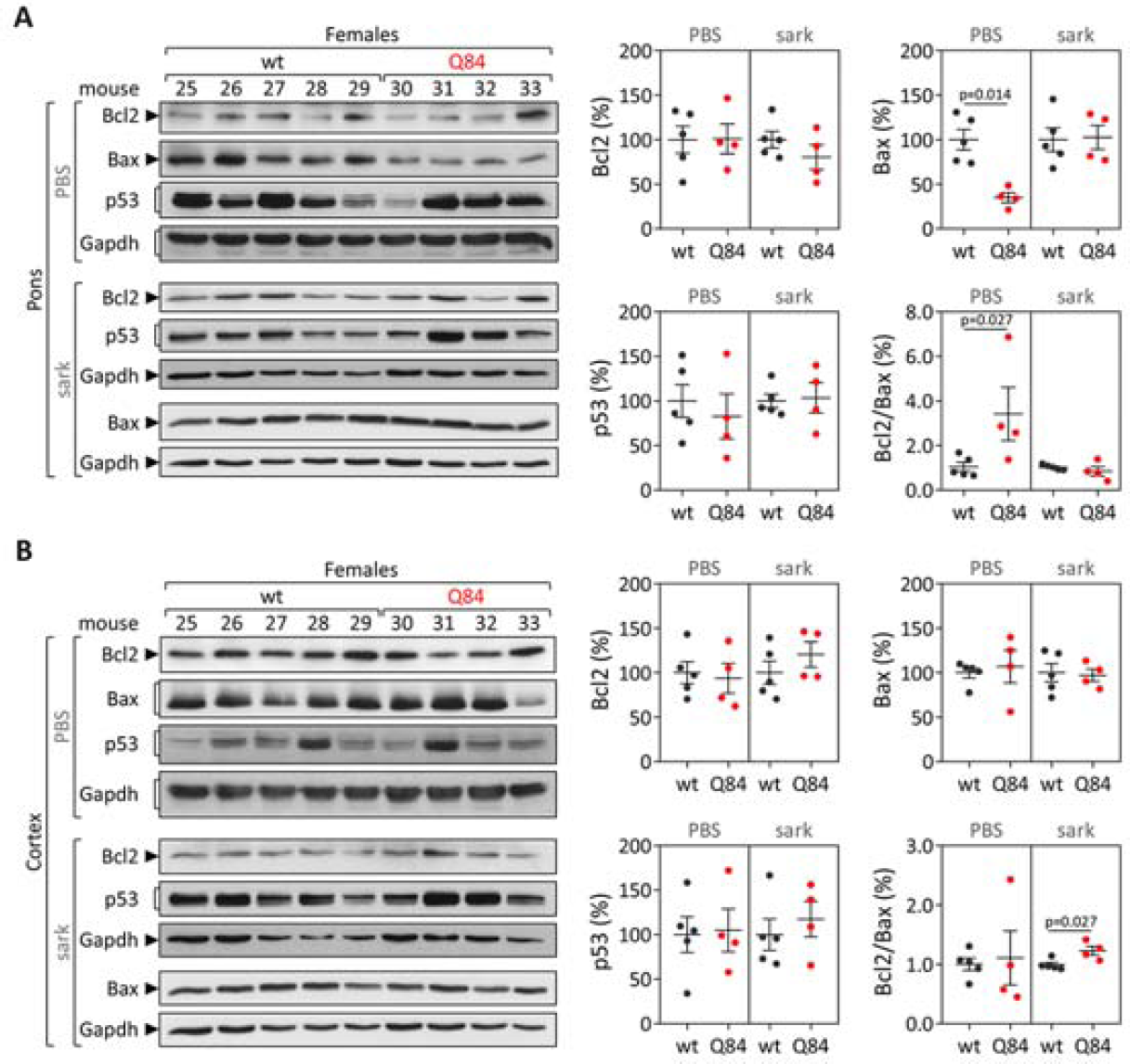
Protein levels of Bcl2, Bax and p53 as well as Bcl2/Bax ratio in PBS-soluble (soluble) and sarkosyl-soluble (insoluble) fractions were assessed in (A) pons and (B) cerebral cortex of symptomatic 18 MO Q84 transgenic and wt mice. Left panels show immunoblots of indicated proteins in both soluble (PBS) and insoluble (sark) fractions. Right panels display quantification of band intensity, with values normalized to GAPDH. Bars in the graphs represent the average percentage of protein relative to controls (± SEM).

## 4. Discussion

In this study the expression behavior of the apoptosis-related genes *BCL2, BAX* and *TP53* and the *BCL2*/*BAX* ratio, an indicator of susceptibility to apoptosis [34], was evaluated in peripheral blood and *post-mortem* brain samples from MJD subjects and Q84 transgenic mice. Like previous reports [8,18], our cross-sectional analysis showed evidence of reduced *BCL2* transcript levels in blood of MJD patients. However, the abundance of blood *BCL2* transcripts revealed a low capacity to discriminate these two biological groups, thus precluding the use of *BCL2* levels as a biomarker of MJD. Similar blood *BCL2*/*BAX* transcript ratio was further found in MJD patients and matched controls, which contrasts with the previous report describing reduced *BCL2*/*BAX* transcript ratio in MJD patients [8]. The partial replication of the published results [8] could perhaps be explained by the analysis of a small number of patients (n=37) and the use of age- and sex-matched paired controls in this study comparing with the study by Raposo and colleagues describing the analysis of a higher number of patients (n=74) but non-matched controls [8]. In addition, similar blood abundance of *BCL2* transcripts and *BCL2*/*BAX* transcript ratio was found here in MJD preclinical subjects and matched controls (n=19), also contrasting with previous reports showing reduced *BCL2* levels [18] and *BCL2*/*BAX* transcript ratio in blood of preclinical subjects (n=16) [8]. Again, the different observations in MJD preclinical subjects here and in the previous reports [8,18] could be due to the use of different experimental designs, namely the use of age- and sex-matched paired controls in this study versus non-matched controls used in the published reports [8,18]. Noteworthy, in this study increased levels of blood pro-apoptotic *BAX* transcripts and decreased *BCL2*/*BAX* ratio were found to be associated with earlier age at onset, indicating that these transcriptional alterations may be associated with MJD pathogenesis. Consistently with these findings, and although global *TP53* levels are similar in patients and healthy controls, our follow-up analysis of blood samples from 18 MJD patients collected at distinct moments of disease progression over a maximum period of nine years showed that the abundance of pro-apoptotic *TP53* transcripts increases with disease progression. In contrast, and while in a lower extent than *TP53* transcript changes, levels of anti-apoptotic *BCL2* transcripts also increased with disease progression. The increased abundance of blood *TP53* and *BCL2* transcripts with MJD progression may indicate that pro-apoptotic and anti-apoptotic signs increase concomitantly in this tissue with the disease course.

While *post-mortem* brains from MJD patients are very rare, dispersed throughout several brain banks in the world and studies using such materials usually show a low power, the analysis of human MJD brain samples contributes to the elucidation of altered cellular mechanisms at the end stage of disease that may be associated with MJD pathogenesis. Overall, our cross-sectional findings of increased *BCL2*/*BAX* transcript ratio in DCN and increased insoluble protein BCL2/BAX ratio in DCN and pons of *post-mortem* MJD brains suggest that, contrarily to what would be expected, cells in the DCN and pons, which are brain areas severely affected by degeneration in MJD [4,28,29] might be more prone to survive compared with controls. Interestingly, this behavior was not observed in the less-affected frontal cortex of MJD patients that only showed significantly increased soluble BAX levels, suggesting that this brain region displays similar susceptibility to apoptosis in MJD patients and in control individuals. The evidence of increased abundance of insoluble BCL2/BAX ratio observed in the DCN and pons of MJD patients may indicate increased abundance of the BCL2 protein in the mitochondrial outer membrane thus preventing apoptosis activation by inhibiting the activation of the pro-apoptotic BCL2 family members [35]. However, the possibility that some of these proteins could instead be recruited to ATXN3 aggregates or be localized in other cellular membranes cannot be excluded. Yet, the results observed in the DCN and pons of MJD patients agree with previous reports of absence of TUNEL-positive cells and changes of expression of apoptosis-related proteins (BCL2, P53, BAX or CPP32) in *post-mortem* DCN from MJD patients [24]. MJD patients globally show a long survival time (time elapsed from onset to death) [36] and thus *post-mortem* MJD brain samples most likely represent end stages of the disease. Hence, the most affected neurons and other cells in the DCN and pons, severely affected with cell loss in MJD patients [4], are most probably dead and absent at the end stage of disease and, therefore, the increased anti-apoptotic signs (BCL2/BAX ratio) observed in both brain regions could indicate that the surviving glial and neuronal cells in these two affected regions activate survival protection mechanisms at the end-stage of MJD. Similarly, Satou et al. [37] reported that the abundance of BCL2 protein within neurons of *post-mortem* brains from Alzheimer’s disease patients increased with disease severity, suggesting that BCL2 protein may have a protective role at the end-stage of the disease.

In parallel with the study in human subjects, mouse *Bcl2, Bax* and *Trp53* transcript and protein levels were assessed in blood of pre-symptomatic 9 MO Q84 transgenic mice, and in pons and cerebral cortex of both pre-symptomatic 9 MO and symptomatic 18 MO Q84 mice [10,30] to evaluate whether this widely used MJD mouse model replicates the findings observed in MJD subjects. Our cross-sectional analysis of Q84 mice, showed similar *Bcl2, Bax* and *Trp53* transcript levels in blood of pre-symptomatic 9 MO Q84 mice and controls. These results in the pre-symptomatic Q84 mice agree with the observed lack of differences between the abundance of *BCL2, BAX* and *TP53* transcripts in peripheral blood samples from preclinical MJD subjects and age- and sex-matched paired controls. Additionally, similar transcript levels of *Bcl2, Bax* and *Trp53* were found in pons and cerebral cortex of pre-symptomatic 9 MO or symptomatic 18 MO Q84 transgenic mice and wt controls. Interestingly, the similar abundance of the three assessed transcripts in pons and cerebral cortex of symptomatic Q84 mice and controls mimics the observations in *post-mortem* brains from MJD patients further indicating that the Q84 mice replicate several aspects of the human disease. In contrast to the observed similar abundance of *Bcl2, Bax* and *Trp53* transcripts in pre-symptomatic and symptomatic Q84 and wt mice, altered levels of Bcl2 and/or Bax proteins were found in the two assessed brain regions of pre-symptomatic and symptomatic Q84 transgenic mice compared to wt littermates. Despite the altered abundance of these two proteins in pons and cerebral cortex of pre-symptomatic 9 MO Q84 mice, the cells present in these brain regions show similar susceptibility to apoptosis (as shown by the similar insoluble Bcl2/Bax ratio) as in controls. Likewise, cells in pons of symptomatic 18 MO Q84 transgenic mice showed a similar susceptibility to apoptosis compared with controls, while the cells in the less-affected cerebral cortex seem to be more prone to survive displaying significantly increased levels of insoluble Bcl2/Bax ratio. The results found in pre-symptomatic and symptomatic Q84 transgenic mice indicate that Bcl2 and Bax protein expression changed with the disease progression in both brain regions. Additionally, the similar results observed in the less-affected cerebral cortex of symptomatic 18 MO Q84 mice and in the severely affected DCN and pons of MJD patients suggest that symptomatic Q84 mice are at different stages of the disease compared to *post-mortem* MJD brains. Globally, the mitochondrial apoptosis mechanisms in cells present in the highly affected and less-affected brain regions seem to respond differently in an age-dependent way to the damage elicited by the expanded *ATXN3* CAG repeat in MJD.

Overall, our study displays some limitations as the limited statistical power of the analysis of *post-mortem* brain samples due to the small sample size, and the lack of blood transcript abundance behavior over disease progression in Q84 transgenic mice due to the lack of peripheral blood samples from the symptomatic 18 MO transgenic Q84 mice. The fact that no transcriptional differences were found in the Q84 mouse model suggests that this animal model may replicate the human disease, yet future studies using mouse samples from several points along the progression of the disease would need to be performed to test this hypothesis. Importantly, future evaluation of the abundance of a wider number of apoptotic-related molecules including markers of the late phase of apoptosis in peripheral blood samples from MJD subjects and *post-mortem* MJD brains could reveal novel biomarkers of MJD and could help elucidating the apoptotic signaling cascades that are majorly involved in MJD. In addition, because mutant ATXN3 stabilizes P53 leading to increased abundance of p53 in MJD cellular models [21], future quantification of the P53 protein in peripheral blood from MJD carriers could reveal a new progression biomarker for MJD.

## 5. Conclusions

Globally, our findings indicate that there is tissue-specific vulnerability to apoptosis in MJD subjects and that this tissue-dependent behavior is only partially replicated by the MJD Q84 mouse model.

## Supplementary Material

Table S1: Characterization of the MJD subjects (preclinical subjects and patients) and control individuals used in this study (n=124), Table S2: Demographic, genetic, and clinical data of the 18 MJD patients used in the follow-up study, Table S3: Characterization of *post-mortem* human brain samples from MJD patients and controls individuals, and RNA integrity number of each brain samples used in this study, Table S4: Genotypes of the 9 and 18 month-old mice used in this study, Table S5: Correlations between transcript levels of *BCL2, BAX* and *TP53*, and *BCL2*/*BAX* ratio, and demographic, genetic and clinical data of MJD subjects (preclinical individuals and patients), Figure S1: *BCL2, BAX* and *TP53* transcriptional levels and *BCL2*/*BAX* ratio changes over time in the 18 MJD patients analyzed in the follow-up study, Figure S2: Western blot using the anti-ATXN3 antibody (1H9) to detect the native human ATXN3 (nATXN3) and expanded human ATXN3 (eATXN3) in insoluble protein fraction of *post-mortem* human samples from dentate cerebellar nucleus (DCN), pons and frontal cortex (Cortex) of Machado-Joseph disease patients (P) and control subjects (C). GAPDH was used as a protein loading control, Figure S3: Western blot using the anti-ATXN3 antibody (1H9) to detect the mutant human ATXN3 (hATXN3) and endogenous mouse ATXN3 (eATXN3) in soluble fraction protein from (A) cerebral cortex of 9 months-old and (B) 18 months-old hemizygous YACMJD84.2 (Q84) transgenic and wild-type (wt) littermate mice. GAPDH was used as a protein loading control.

## Author Contributions

Conceptualization, M.R, M.C.C. and M.L.; Methodology, A.F.F., M.R., M.C.C., and M.L.; Validation, A.F.F., M.C.C. and M.L.; Formal Analysis, A.F.F., M.F.B., M.C.C., and M.L.; Investigation, A.F.F., M.R., E.D.S., N.S.A., F.M., M.F.B., M.P., J.V., T.K., M.C.C. and M.L.; Resources, M.C.C. and M.L.; Writing – Original Draft Preparation, A.F.F., M.C.C. and M.L.; Writing – Review & Editing, A.F.F., M.R., E.D.S., N.S.A., F.M., M.F.B., M.P., J.V., T.K., M.C.C. and M.L.; Visualization, A.F.F., M.C.C. and M.L.; Supervision, M.C.C. and M.L.; Project Administration, M.C.C. and M.L.; Funding Acquisition, M.C.C. and M.L. All authors have read and agreed to the published version of the manuscript.

## Funding

This work was funded by FEDER funds through the Operational Competitiveness Program (COMPETE) and by National Funds through Fundação para a Ciência e a Tecnologia (FCT) under the project FCOMP-01-0124-FEDER-028753 (PTDC/DTP/PIC/ 0370/2012) and was also funded by National Ataxia Foundation through a research seed money grant (2018). AFF was supported by PhD grant (SFRH/BD/121101/2016) funded by FCT, República Portuguesa/ Ciência, Tecnologia e Ensino Superior, Portugal 2020, União Europeia/Fundo Social Europeu and POR_NORTE. MR is supported by FCT (CEECIND/03018/2018).

## Institutional Review Board Statement

The study was conducted according to the guidelines of the Declaration of Helsinki. Blood samples from MJD subjects and controls were collected in the scope of the projects MESCA3 (PTDC/DTP-PIC/0370/2012), approved by the Ethical Committee of the Hospital do Divino Espírito Santo. The use of *post-mortem* human brain samples was approved by the University of Michigan Biosafety Committee (IBCA00000265, 7th March 2017). Animal procedures were approved by the University of Michigan Committee on Use and Care of Animals (Protocol PRO00006371, 6th October 2015). Project research seed money grant “Apoptosis-related genes BCL2, BAX and TP53 as biomarkers of Machado-Joseph Disease”, approved by the Ethical Committee of the University of the Azores (PARECER 26/2018, 25th September 2018) and by the University of Michigan (AWD006601, 9th January 2018).

## Informed Consent Statement

Informed consent was obtained from all subjects involved in the study.

## Data Availability Statement

The data presented in this study are available on request from the corresponding author.

## Acknowledgments

We are grateful to the MJD subjects and their relatives as well as to the healthy volunteers for participating in this study. We thank Ilya Bezprozvanny for sharing the YACMJD84.2 mice. The Michigan brain bank at the University of Michigan is partially supported by the NIH/NIA funded Michigan Alzheimer’s Disease Center (P30AG053760 and P30AG072931). The Centro de Estatística e Aplicações is partially supported by National Funds through Fundação para a Ciência e a Tecnologia (FCT), Portugal, project UIDB/MAT/ 00006/2020 (CEA/UL).

## Conflicts of Interest

The authors declare no conflict of interest.

## Notes

### Competing Interest Statement

The authors have declared no competing interest.

