## Supplementary material for "Tissue-specific vulnerability to apoptosis in Machado-Joseph disease"

Supplementary Table S1. Characterization of the MJD subjects (preclinical subjects and patients) and control individuals used in this study (n=124).

| | **Preclinical subjects** | **Patients** | **Controls** |
|---|---|---|---|
| **Blood samples** | | | |
| n (Female; Male) | 19 (12; 7) | 37 (19; 18) | 54[†] (30; 24) |
| Age[1], years | 30.3 ± 7.4 [21; 44] | 45.9 ± 11.6 [26; 65] | 41.1 ± 12.7 [21; 67] |
| $CAG_n$ allele 1[2] | 20.0 ± 4.1 [14; 28] | 20.7 ± 4.9 [14; 29] | 19.6 ± 4.3 [14; 27]* |
| $CAG_n$ allele 2[3] | 68.1 ± 3.0 [62; 75] | 71.2 ± 2.8 [64; 76] | 24.0 ± 4.0 [14; 32]* |
| Years to onset[4], years | -10.5 ± 9.2 [-26; +5] | NA | NA |
| Age at onset, years | NA | 35.2 ± 7.9 [22; 52] | NA |
| Disease duration, years | NA | 10.7 ± 8.7 [1; 36] | NA |
| ***Post-mortem* brain samples** | | | |
| n (Female; Male) | NA | 5 (4; 1) | 9 (5; 4) |
| $CAG_n$ allele 1[2] | NA | 19.0 ± 4.1 [12; 22] | 17.0 ± 5.3 [11; 25] |
| $CAG_n$ allele 2[3] | NA | 70.2 ± 3.0 [66; 73] | 22.6 ± 5.6 [12; 30] |
| Age at onset, years | NA | 45.0 ± 8.5 [39; 51]** | NA |
| Disease duration, years | NA | 26.5 ± 9.2 [20; 33]** | NA |
| Age at death, years | NA | 63.0 ± 16.0 [48; 84] | 69.6 ± 12.2 [48; 83] |
| PMI[5], hours | NA | 26.6 ± 17.2 [4; 48] | 15.4 ± 8.0 [4; 24] |

Quantitative variables are displayed as mean ± standard deviation [minimum; maximum]; [1]Age at first blood collection, [2]number of CAG repeats in the normal allele of MJD subjects/number of CAG repeats in normal allele 1 of controls; [3]number of CAG repeats in expanded allele of MJD subjects/number of CAG repeats in normal allele 2 of controls; [4]Years to onset: negative values indicate the numbers of years missing to the estimated onset and positive values indicate the number of years that have elapsed the estimated onset; [5]*Post-mortem* interval; [†]Age (±3 years) and sex-matched paired controls for preclinical subjects and patients (two individuals were used as paired matched controls for both groups); *Information available for 47 controls; **Information available for two patients; NA, not applicable/not available

Supplementary Table S2. Demographic, genetic, and clinical data of the 18 MJD patients used in the follow-up study.

| | **Baseline** | **Visit 1** | **Visit 2** |
|---|---|---|---|
| n (Female; Male) | 18 (7; 11) | 18 (7; 11) | 11 (4; 7) |
| Age[1], years | 48.9 ± 13.7 [26; 65] | 53.9 ± 13.6 [32; 72] | 52.6 ± 13.9 [34; 72] |
| Normal CAG allele | 19.7 ± 5.1 [14; 29] | 19.7 ± 5.1 [14; 29] | 19.6 ± 5.9 [14; 29] |
| Expanded CAG allele | 70.9 ± 3.2 [64; 76] | 70.9 ± 3.2 [64; 76] | 71.4 ± 3.9 [64; 76] |
| Age at onset, years | 36.2 ± 7.9 [22; 50] | 36.2 ± 7.9 [22; 50] | 36.1 ± 9.0 [22; 50] |
| Disease duration, years | 12.7 ± 9.5 [1; 36] | 17.7 ± 9.4 [7; 40] | 16.6 ± 6.7 [9; 27] |

Quantitative variables are displayed as mean ± standard deviation [minimum; maximum]; [1]Age at first blood collection

Supplementary Table S3. Characterization of *post-mortem* human brain samples from MJD patients and controls individuals, and RNA integrity number of each brain samples used in this study.

| Health condition | Sex | ID | Age at death (years) | PMI[1] (hours) | Cause of death | Age at onset (years) | CAG repeats | RNA integrity number | | |
|---|---|---|---|---|---|---|---|---|---|---|
| | | | | | | | Allele 1 \| Allele 2 | DCN[2] | Pons | Frontal cortex |
| Controls | Female | 1 | 48 | 5 | Polycythemia vera, mesenteric thyrombosis and ischemic bowel resection | NA | 12\|19 | 3.7 | 3.2 | 3.1 |
| | | 2 | 76 | 14 | Cardiac failure | NA | 21\|25 | 2.9 | 5.3 | 3.4 |
| | | 3 | 80 | 19 | Congestive heart failure and atrial fibrillation | NA | 12\|18 | 5.2 | 6 | NA |
| | | 4 | 83 | NA | Renal cell carcinoma | NA | 18\|21 | 4.8 | 4.6 | 5.6 |
| | | 5 | 83 | 21 | Cardiac arrest, urinary tract infection and sepsis | NA | 11\|12 | 6.9 | 5.7 | NA |
| | Male | 6 | 59 | 12 | Sudden cardiac arrest, ventricular fibrillation and post-shock electromechanical dissociation | NA | 12\|25 | 7.7 | NA | 6.5 |
| | | 7 | 61 | 24 | Cardiac failure, cardiogenic shock and post-shock electromechanical dissociation | NA | 21\|25 | 7.5 | NA | 6.9 |
| | | 8 | 65 | 24 | Acute respiratory distress syndrome and sepsis | NA | 25\|28 | NA | NA | NA |
| | | 9 | 71 | 4 | Cardiac failure | NA | 21\|30 | 7 | 7.4 | 8.3 |
| MJD patients | Female | 10 | 48 | 22 | NA | NA | 22\|73 | 5.3 | 4.1 | 6.2 |
| | | 11 | 59 | 4 | NA | 39 | 21\|70 | 7.9 | 6.2 | 8.4 |
| | | 12 | 75 | 39 | NA | NA | 19\|69 | NA | NA | 4 |
| | | 13 | 84 | 20 | NA | 51 | 21\|66 | 6.5 | NA | 4.7 |
| | Male | 14 | 49 | 48 | NA | NA | 12\|73 | NA | 3.5 | 3.9 |

[1]*Post-mortem* interval; [2]Dentate cerebellar nucleus; NA, not applicable/not available

Supplementary Table S4. Genotypes of the 9 and 18 month-old mice used in this study.

| Age | Genotype | Gender | Mouse # | Mouse Tag | CAG repeats |
|---|---|---|---|---|---|
| 9-month-old | wt[1] | Female | 1 | 840.0.0 | NA |
| | | | 2 | 840.0.2 | |
| | | | 3 | 817.0.0 | |
| | | | 4 | 817.0.3 | |
| | | | 5 | 843.0.1 | |
| | | | 6# | 840.0.4 | |
| | | Male | 7 | 842.0.1 | NA |
| | | | 8 | 845.0.1 | |
| | | | 9 | 845.0.2 | |
| | | | 10 | 839.0.2 | |
| | | | 11* | 846.0.2 | |
| | | | 12* | 855.0.0 | |
| | Q84[2] | Female | 13 | 817.0.1 | 73/74/**76**/83 |
| | | | 14 | 817.0.2 | **73**/76/79/88 |
| | | | 15 | 840.0.1 | **72**/75/84 |
| | | | 16 | 817.0.4 | 68/**73**/80/83 |
| | | | 17 | 843.0.0 | 68/**73**/79/82 |
| | | | 18 | 843.0.3 | **71**/75/78/85 |
| | | Male | 19 | 813.0.0 | 68/**73**/77/82 |
| | | | 20 | 813.0.1 | 68/**73**/77/83 |
| | | | 21 | 842.0.2 | **71**/75/81 |
| | | | 22 | 845.0.0 | **72**/76/86 |
| | | | 23* | 845.0.3 | **72**/77/85 |
| | | | 24# | 842.0.0 | NA |
| 18-month-old | wt[1] | Female | 25 | 61.0.0 | NA |
| | | | 26 | 61.0.1 | |
| | | | 27 | 691.0.3 | |
| | | | 28 | 702.0.0 | |
| | | | 29 | 702.0.1 | |
| | Q84[2] | | 30 | 61.0. 4 | NA |
| | | | 31 | 691.0.1 | |
| | | | 32* | 702.0.2 | |
| | | | 33 | 710.0.3 | |

[1]wild type littermate mice; [2]hemizygous YACMJD84.2 transgenic mice; #Protein sample not available; *RNA sample not available; NA, not available; the main CAG allele is indicated in bold

Supplementary Table S5. Correlations between transcript levels of *BCL2*, *BAX* and *TP53*, as well as *BCL2*/*BAX* ratio and demographic, genetic and clinical data of MJD subjects (preclinical individuals and patients).

| | | **Age[1]** | **CAG-E[2]** | **Years to onset** | **Age at onset** | | **DD[3]** | |
|---|---|---|---|---|---|---|---|---|
| | | | | | **CAG-E adj.** | **Age + CAG-E adj.** | **No adj.** | **Age[1] adj.** |
| **Preclinical subject** | | | | | | | | |
| *BCL2* | Rho | 0.269 | 0.292 | -0.275 | NA | NA | NA | NA |
| | Sig (2-tailed) | 0.266 | 0.225 | 0.255 | | | | |
| *BAX* | Rho | -0.001 | 0.052 | 0.002 | NA | NA | NA | NA |
| | Sig (2-tailed) | 0.997 | 0.832 | 0.994 | | | | |
| *TP53* | Rho | -0.090 | 0.078 | 0.101 | NA | NA | NA | NA |
| | Sig (2-tailed) | 0.715 | 0.751 | 0.681 | | | | |
| *BCL2*/*BAX* | Rho | 0.005 | 0.217 | 0.093 | NA | NA | NA | NA |
| | Sig (2-tailed) | 0.985 | 0.403 | 0.722 | | | | |
| **Patient** | | | | | | | | |
| *BCL2* | Rho | -0.264 | 0.268 | NA | 0.011 | NA | 0.229 | NA |
| | Sig (2-tailed) | 0.114 | 0.109 | | 0.951 | | 0.173 | |
| *BAX* | Rho | -0.350 | 0.273 | NA | -0.522 | -0.482 | 0.048 | 0.308 |
| | Sig (2-tailed) | **0.034** | 0.102 | | **0.001** | **0.003** | 0.777 | 0.067 |
| *TP53* | Rho | -0.360 | 0.295 | NA | -0.189 | -0.106 | 0.198 | 0.090 |
| | Sig (2-tailed) | **0.029** | 0.076 | | 0.270 | 0.544 | 0.240 | 0.600 |
| *BCL2*/*BAX* | Rho | 0.243 | -0.222 | NA | 0.403 | 0.393 | 0.014 | 0.279 |
| | Sig (2-tailed) | 0.153 | 0.194 | | **0.016** | **0.022** | 0.933 | 0.105 |

[1]Age at first blood collection; [2]CAG-E: expanded CAG repeat;[3]Disease duration

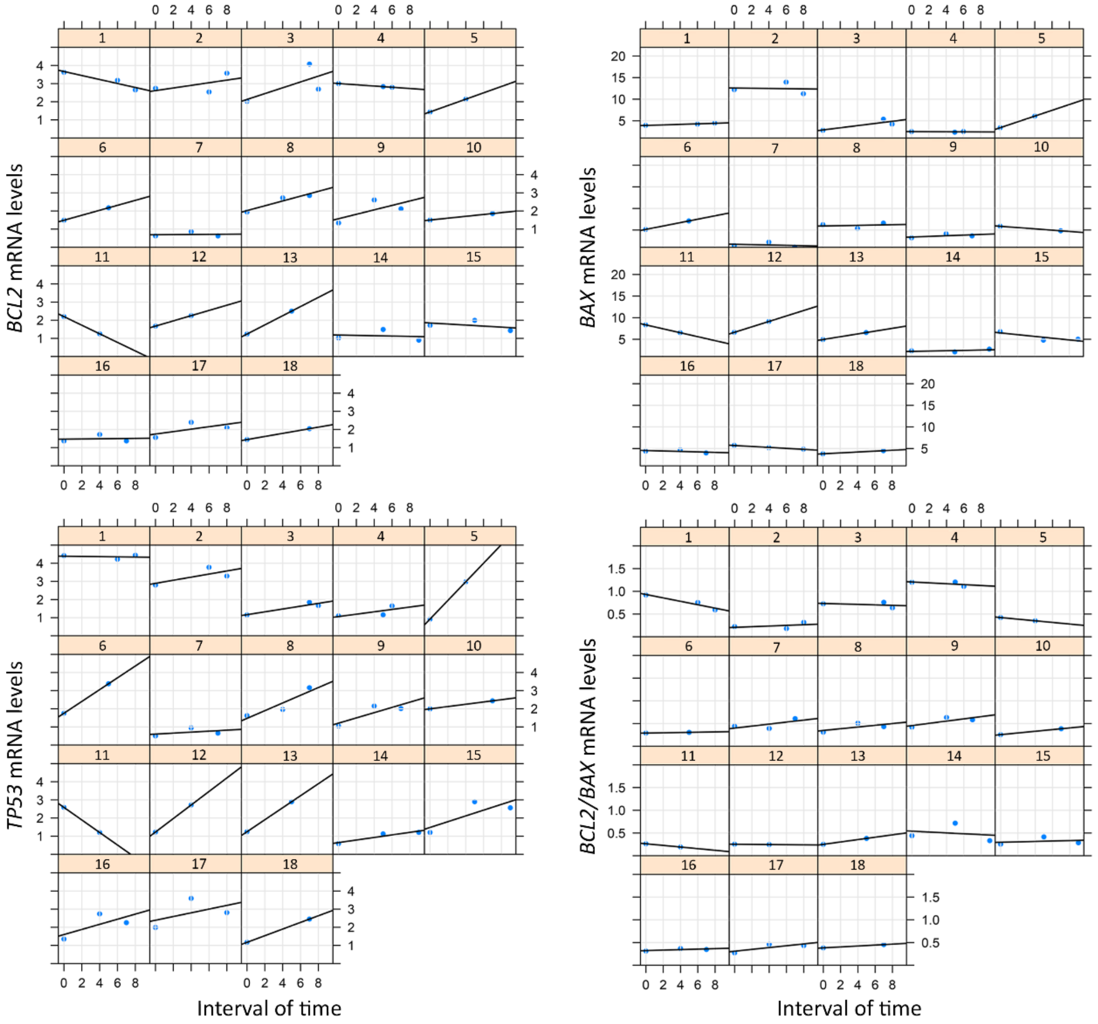


Supplementary Figure S1. *BCL2*, *BAX* and *TP53* transcriptional levels and *BCL2*/*BAX* ratio changes over time in the 18 patients analyzed in the follow-up study.

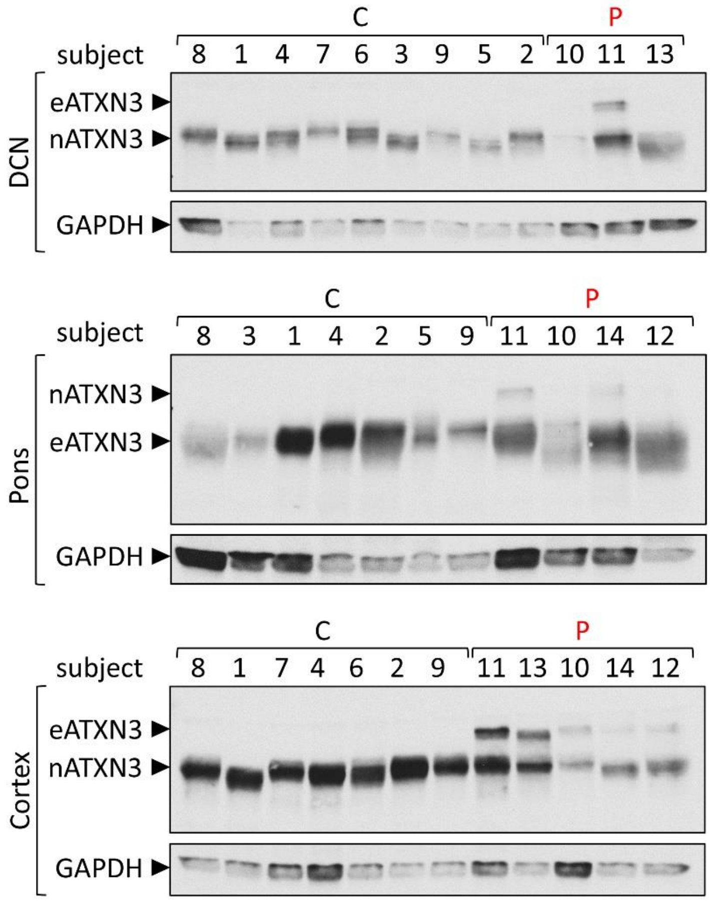


Supplementary Figure S2. Western blot using the anti-ATXN3 antibody (1H9) to detect the native human ATXN3 (nATXN3) and expanded human ATXN3 (eATXN3) in insoluble protein fraction of *post-mortem* human samples from dentate cerebellar nucleus (DCN), pons and frontal cortex (Cortex) of Machado-Joseph disease patients (P) and control subjects (C). GAPDH was used as a protein loading control.

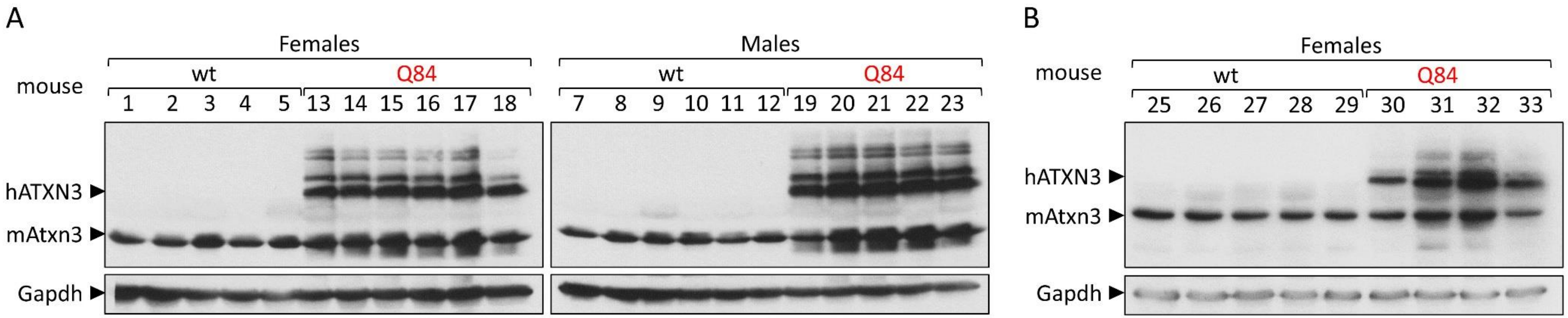


Supplementary Figure S3. Western blot using the anti-ATXN3 antibody (1H9) to detect the mutant human ATXN3 (hATXN3) and endogenous mouse ATXN3 (eATXN3) in soluble fraction protein from (A) cerebral cortex of 9 months-old and (B) 18 months-old hemizygous YACMJD84.2 (Q84) transgenic and wild-type (wt) littermate mice. GAPDH was used as a protein loading control.
